# Effects of different shading levels on growth, physiology and leaf surface micromorphology of *Sorbus sibirica* ‘Dong Hong’ seedlings

**DOI:** 10.64898/2026.07.25.740310

**Authors:** Rongyan Huang, Wenhao Gong, Xinyuan Li, Shida Ji, Tingting Cui, Lanlan Zhang

## Abstract

**Background:** Shading may influence nursery cultivation of *Sorbus sibirica* ‘Dong Hong’, but its growth and physiological responses remain insufficiently characterised.

**Aims:** To assess seedling responses to shading and inform container nursery management.

**Methods:** One-year-old potted seedlings were grown under full sunlight (CK) and nominal shading levels of 30% (L1), 50% (L2) and 70% (L3) for 100 days in Qiqihar, China. Growth, biomass allocation, root morphology, physiological traits and leaf micromorphology were assessed.

**Results:** Seedling height, total biomass, chlorophyll and soluble protein contents peaked under L1, whereas basal diameter and invertase activity declined with increasing shade. Under L1, total root length and mean root diameter were comparable to CK, but root tip number and root volume were lower. Root biomass proportion remained similar across treatments. L2 and L3 had higher leaf biomass proportions but lower total biomass than CK. Shaded seedlings had smaller stomatal dimensions. Biomass and root morphological values were lowest under L3.

**Conclusions:** L1 combined greater biomass with thinner stems and altered root morphology. Restricted rooting volume, limited environmental monitoring and a single summer-to-autumn trial warrant validation before extending these findings to other seasons, regions or cultivation systems.

**Highlights:**

- 30% shade increased biomass of *Sorbus sibirica* ‘Dong Hong’ seedlings.
- 30% shade increased chlorophyll and soluble protein but reduced stem thickness.
- 70% shade yielded the lowest biomass and root morphological values.

## Introduction

Light availability is an important factor in nursery management because seedling morphology, growth and biomass allocation can vary with light conditions (Sevillano et al., 2016). Shading can modify the light environment, but its effects depend on the balance between reducing excessive radiation and maintaining sufficient light for carbon assimilation (Jiang et al., 2026). Intermediate shading has been associated with favourable growth and physiological responses in some nursery species, whereas stronger shading may limit biomass production (Li et al., 2025). However, different traits need not respond consistently: greater seedling height (SH) may coexist with reduced stem thickness or changes in root development. Evaluating shading practices therefore requires consideration of several traits rather than reliance on height or leaf colour alone (Alagupalamuthirsolai et al., 2018; Cheng et al., 2020).

*Sorbus sibirica* ‘Dong Hong’ is a deciduous tree in the family Rosaceae, reaching 6–12 m in height. It was selected by the Heihe Academy of Forestry Sciences, China, from *S. sibirica* germplasm introduced from a botanical garden in Siberia, Russia, and has been described as suitable for urban landscaping in northern China (Xie et al., 2017). Its reported ornamental features include red autumn foliage and red to orange-red fruits retained into winter, providing seasonal interest in cold-region landscapes (Cao et al., 2019). In a comparative study with *S. pohuashanensis*, another congener occurring in Northeast China, ‘Dong Hong’ exhibited greater mean plant height and basal diameter, significantly higher seed-setting and fruit-setting rates per inflorescence, and a 100-fruit mass 1.45 times that of *S. pohuashanensis* (Chai et al., 2015; Yin et al., 2017). Although both taxa produce red or orange-red fruits, these reported differences in growth and reproductive traits support further investigation of ‘Dong Hong’ for landscape use (Liu et al., 2022; Zhao, 2022). Its ornamental characteristics and potential applications highlight the need to establish suitable nursery practices, while its comparative market value and environmental tolerance require further evaluation.

Previous work has examined the responses of ‘Dong Hong’ to saline–alkali stress, providing information on its environmental sensitivity and physiological adjustment (Huang et al., 2024). Nevertheless, responses to soil stress do not establish an appropriate nursery light regime. For container-grown seedlings, light management also interacts with restricted rooting space and the availability of water and nutrients (Chen et al., 2025a). Strong radiation may increase thermal and evaporative demands, whereas prolonged or excessive shading may restrict growth (Qin et al., 2022). These represent potential management risks rather than demonstrated causes of poor performance in local ‘Dong Hong’ nurseries. A practical question is whether shading can improve useful seedling traits without compromising stem development and root characteristics. Selecting shade nets solely by nominal shading rate or apparent shoot elongation may overlook these differences (Qin et al., 2024; Li et al., 2025). Evidence from related species indicates that shading responses within Sorbus warrant investigation but cannot be transferred directly between taxa. Research reported favourable growth and photosynthetic performance in *S. tianschanica* seedlings at approximately 30% of full sunlight (Chen et al., 2025b). Their findings indicate that morphological and physiological adjustment can contribute to light responses within Sorbus, but cannot directly establish an appropriate shading regime for ‘Dong Hong’.

Several processes could explain favourable responses of individual traits to shading. Reduced radiation may lessen excitation pressure or thermal load, while changes in pigment content and shoot development may improve light capture (Bashyal et al., 2025; Patil and Deshmukh, 2026). These adjustments need not increase whole-plant biomass or improve all aspects of seedling structure (Ushimaru et al., 2025). Changes in root morphology and leaf surface traits alone cannot establish improvements in resource uptake or water-use efficiency (Inurreta-Aguirre et al., 2022; Huang et al., 2025). The unresolved question for ‘Dong Hong’ is therefore whether responses in height and physiological traits coincide with changes in biomass, stem thickness and root development. Addressing this question can help distinguish improvements in selected traits from broader benefits for nursery production (Isaac et al., 2024; Mensah et al., 2024).

This study compared one-year-old potted *S. sibirica* ‘Dong Hong’ seedlings under full sunlight (CK) and nominal shading levels of 30% (L1), 50% (L2) and 70% (L3) during a 100-day summer-to-autumn experiment in Qiqihar, Northeast China. We assessed shoot growth, biomass allocation, root morphology, chlorophyll (CHL) content and soluble protein (SP) content, invertase (INV) activity and leaf surface micromorphology. The objectives were to characterise trait-specific responses, determine whether greater height coincided with greater biomass and stem development, and examine associations among growth, physiological and morphological traits. The study was intended to inform the evaluation of shading options for container nurseries, while recognising that a single cultivation period cannot establish a universally optimal regime across seasons, regions or field planting conditions.

## Materials and methods

### Experimental materials and shading treatments

One-year-old seedlings of *Sorbus sibirica* ‘Dong Hong’ were obtained from Qiqihar Branch of the Heilongjiang Academy of Forestry Sciences. Healthy seedlings of similar initial size were transplanted on 20 June 2024 at the teaching and training base of Qiqihar University, Qiqihar, Heilongjiang Province, China (47.35°N, 123.92°E). Each plastic nursery pot contained one seedling and measured 15 cm in bottom diameter, 21 cm in top diameter and 17.5 cm in height. The substrate consisted of perlite and garden soil mixed at 1:2 (v/v). Four treatments were established: full sunlight (CK) and nominal shading levels of 30% (L1), 50% (L2) and 70% (L3). The shaded treatments used single layers of 2-, 3- and 6-needle black shade netting, respectively. Treatments began on 20 July 2024 and ended on 28 October 2024, corresponding to an elapsed duration of 100 days. The experiment included 50 pots per treatment and 200 pots in total.

Each treatment was applied within a single structure containing 50 pots, giving four structures in total. Each structure measured 3 × 2.5 × 2 m (length × width × height), and adjacent structures were separated by 3 m. For the shading treatments, shade netting covered the roof and all four sides. The CK treatment used an identical frame without shade netting. Pots remained in fixed positions throughout the treatment period. Seedlings were watered manually every two days, and no fertiliser was applied during treatment. Air temperature and relative humidity were not monitored.

Illuminance was measured at seedling canopy height using a TES-1334A illuminance meter (TES Electrical Electronic Corp., Taiwan, China), with an upper measurement limit of 20 000 lx. Measurements were taken at five randomly selected points within each treatment at 10:00, 12:00, 14:00 and 16:00 on three clear-sky days: 25 July, 10 August and 1 September 2024. The sensor was held horizontally. The five point readings were averaged at each measurement time, and treatment values were summarised as the mean ± standard deviation (SD) of the 12 date–time means. Mean illuminance was 18 520 ± 146 lx under CK, 12 601 ± 232 lx under L1, 9 136 ± 189 lx under L2 and 5 640 ± 377 lx under L3. The resulting attenuation values were approximately 32.0%, 50.7% and 69.5% for L1, L2 and L3, respectively. No over-range readings were recorded. Treatment labels denote nominal shading levels; measured illuminance attenuation is reported separately. Illuminance was not converted to photosynthetic photon flux density. E_s_ and E_o_ represent mean illuminance under the shaded treatment and CK. Relative illuminance attenuation was calculated from the overall treatment means using the following equation (Chen et al., 2025b; Noor et al., 2026):

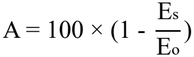

### Measurement of indicators

### Measurement of growth parameters

At the end of the treatment period, five seedlings per treatment were randomly selected for measurements of SH, basal diameter (BD), root morphology and biomass. Growth measurements and root scanning were completed before the same seedlings were separated into organs for dry biomass determination. Physiological assays and scanning electron microscopy (SEM) observations were performed on separately sampled seedlings.

SH was measured from the substrate surface to the shoot apex using a measuring tape. BD was measured approximately 1 cm above the substrate surface using vernier calipers. Measurements represented final seedling dimensions rather than changes from pretreatment values (Chen et al., 2025c).

### Biomass determination

Sampled seedlings were separated into roots, stems including petioles, and leaf blades. Samples were dried at 105 °C for 1 h and subsequently at 75 °C to constant mass. Before weighing, samples were cooled to room temperature in a desiccator. Dry mass was determined using an analytical balance with a readability of 0.0001 g. Constant mass was defined as a difference of no more than 0.002 g between two successive weighings following an additional 2 h of drying at 75 °C.

Leaf biomass (LB), stem biomass (SB) and root biomass (RB) were expressed as dry mass per seedling. Total biomass (TB) was calculated as: TB = LB + SB + RB.

For each seedling, leaf biomass proportion (LBP), stem biomass proportion (SBP) and root biomass proportion (RBP) were calculated as (Read et al., 2017):

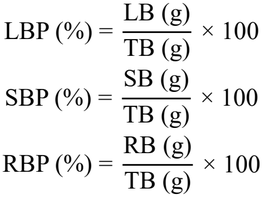

### Determination of root morphology parameters

Root systems from five seedlings per treatment were analysed individually. Roots were carefully washed, and excess surface moisture was removed with absorbent paper. Each root system was spread in a single layer and scanned using an LA-S2400 root scanner (Hangzhou Top Instrument Co., Ltd., Hangzhou, China) at 400 dpi. Images were inspected for overlapping roots; roots were repositioned and rescanned when overlap prevented identification. Images of the complete root system of each sampled seedling were analysed using WinRHIZO Pro 2017 (Regent Instruments Inc., Quebec, Canada) with the default analysis settings. Root surface area (RSA), root tip number (NR), mean root diameter (RAD), root volume (RV) and total root length (TRL) were determined (Lin et al., 2026).

### Measurement of physiological parameters

Five additional seedlings per treatment were randomly selected for physiological measurements. Leaf samples from each seedling were divided into separate portions for CHL content, INV activity and SP content assays. Samples from individual seedlings were kept separate, and three technical determinations per assay were averaged to obtain one value per seedling. Leaf samples were collected between 09:00 and 10:00 at the end of the treatment period. Leaflet blades were sampled from the third to fifth fully expanded compound leaves counted from the shoot apex, with petioles and prominent midribs removed. Sampling was alternated among treatments. All samples were transported on ice in darkness, and fresh tissue was used for CHL content, INV activity and SP content analyses. Extraction for all three assays was completed on the day of sampling. CHL content and SP content were expressed as mg·g^-1^ fresh weight (FW), and INV activity was expressed as U·g^-1^ FW.

For total CHL content determination, 0.2 g of fresh leaf tissue was extracted using 80% (v/v) acetone. Extraction was repeated until the residue was essentially colourless, and the combined extracts were adjusted to 10 mL. Absorbance was measured at 645 and 663 nm in a 1-cm-path-length cuvette, using 80% acetone as the blank (Yang et al., 2026). Total CHL content was calculated as:

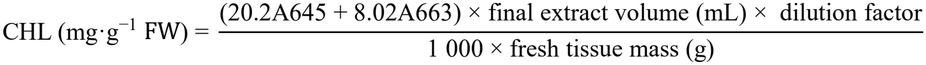

For INV activity determination, 0.2 g of fresh leaf tissue was homogenised in 2.0 mL of 50 mM sodium phosphate buffer (pH 7.0) containing 1 mM ethylenediaminetetraacetic acid (EDTA) and 1% (w/v) polyvinylpolypyrrolidone (PVPP). The homogenate was centrifuged at 12 000 × g for 15 min at 4 °C. The 1.0-mL reaction mixture contained 0.1 mL of supernatant, 100 mM sucrose and 50 mM sodium acetate buffer (pH 4.5), expressed as final concentrations. Reactions were incubated at 37 °C for 30 min. Reducing sugars were measured using the 3,5-dinitrosalicylic acid (DNS) method at 540 nm against a glucose standard curve, with reagent and zero-time controls. One unit was defined as the production of 1 μmol of reducing sugars, expressed as glucose equivalents, per minute under the assay conditions (Chen et al., 2026).

For SP content determination, 0.2 g of fresh leaf tissue was homogenised in 2 mL of 50 mM sodium phosphate buffer (pH 7.0). Following centrifugation at 12 000 × g for 15 min at 4 °C, protein in the supernatant was determined by the Bradford method at 595 nm, using bovine serum albumin as the standard (Bradford, 1976). SP content was calculated from the protein concentration determined using the standard curve, the extraction volume and any dilution factor, and expressed per gram of leaf FW.

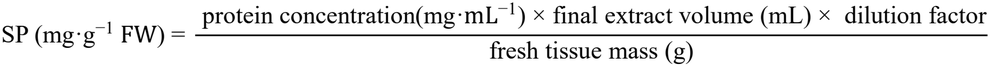

### Observation of leaf surface morphology

A separate set of five seedlings per treatment was randomly selected for SEM. These seedlings were distinct from those used for growth, root morphology, biomass and physiological measurements. The third fully expanded compound leaf counted from the shoot apex was sampled from each seedling. Tissue segments were excised from the middle of the leaflet blade, avoiding the main vein, and the abaxial surface was examined. Samples were fixed in 2.5% glutaraldehyde overnight at 4 °C and rinsed three times with 0.1 M phosphate buffer (pH 7.2) for 15 min each. Samples were then trimmed to approximately 1 × 1 mm and post-fixed in 1% osmium tetroxide for 2 h. After rinsing, samples were dehydrated through an ethanol series of 30%, 50%, 70%, 80%, 90% and 95%, with 15 min at each concentration, followed by three 15-min changes of absolute ethanol. Samples underwent CO_2_ critical-point drying, with final conditions of 35 °C and 8.0 MPa maintained for 10 min before controlled depressurisation. Dried samples were mounted with the surface of interest facing upwards and sputter-coated with gold at 15 mA for 60 s (Kumachova et al., 2021; Li et al., 2024).

Samples were examined using a HITACHI S-3400N scanning electron microscope (Hitachi High-Technologies Corporation, Tokyo, Japan) at an accelerating voltage of 10 kV and a working distance of 10 mm. Magnifications of 500×, 2000× and 5000× were used to assess stomatal density (SDD), stomatal dimensions and epicuticular wax appearance, respectively. Ten non-overlapping fields on the abaxial surface of each sampled leaflet were examined to determine SDD. For stomata-bearing surfaces, 30 intact stomata per leaflet surface were selected across the images to measure stomatal length (SL) and stomatal area (SA). SL was measured along the longitudinal axis of the stomatal complex, and SA was determined by tracing its outer boundary. Measurements were performed using Motic Images Plus 3.1 (Motic China Group Co., Ltd., Xiamen, China), with spatial calibration based on the scale bar in each original image. Measurements were summarised separately for each seedling and leaf abaxial surface before treatment-level summaries were calculated. Epicuticular wax appearance was assessed qualitatively (Huang et al., 2023). SL was expressed in µm and SA in µm^2^. SDD was calculated as:

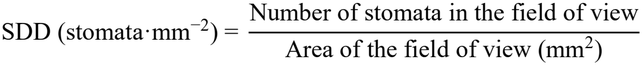

### Data processing

Data were analysed using SPSS 26.0 (IBM Corp., Armonk, NY, USA). Before analysis of variance (ANOVA), residual normality was assessed using the Shapiro–Wilk test and normal quantile-quantile (Q–Q) plots, and homogeneity of variances among treatments was evaluated using Levene’s test. These checks did not indicate substantial departures from the model assumptions; therefore, analyses were performed on untransformed data. Treatment differences were evaluated using one-way ANOVA followed by Tukey’s honestly significant difference test at *P* < 0.05. Seedling-level values were used for these analyses; technical determinations and within-leaf measurements were averaged for each seedling before analysis. Results are presented as means ± SD of five seedlings per treatment, except for illuminance, which was summarised across 12 date–time means.

Spearman rank correlation coefficients (ρ) were calculated from the four treatment means to explore similarities in trait rankings. Given the small sample size (*n* = 4), correlations were interpreted descriptively without significance testing. These associations represent covariation among treatment means, rather than relationships among individual seedlings, and do not establish causality. Figures were prepared using Origin 2021 (OriginLab Corp., Northampton, MA, USA).

## Results

### Shoot growth

SH and BD showed different patterns across shading treatments. SH was highest under L1, followed by L2, CK and L3 (Fig. 1*A*). In contrast, BD decreased progressively from CK to L3 (Fig. 1*B*). Thus, seedlings under L1 and L2 were taller but had thinner stems than those under CK, whereas both traits reached their lowest values under L3. These observations indicate that greater SH under L1 and L2 was not accompanied by greater stem thickness.

**Fig. 1.**
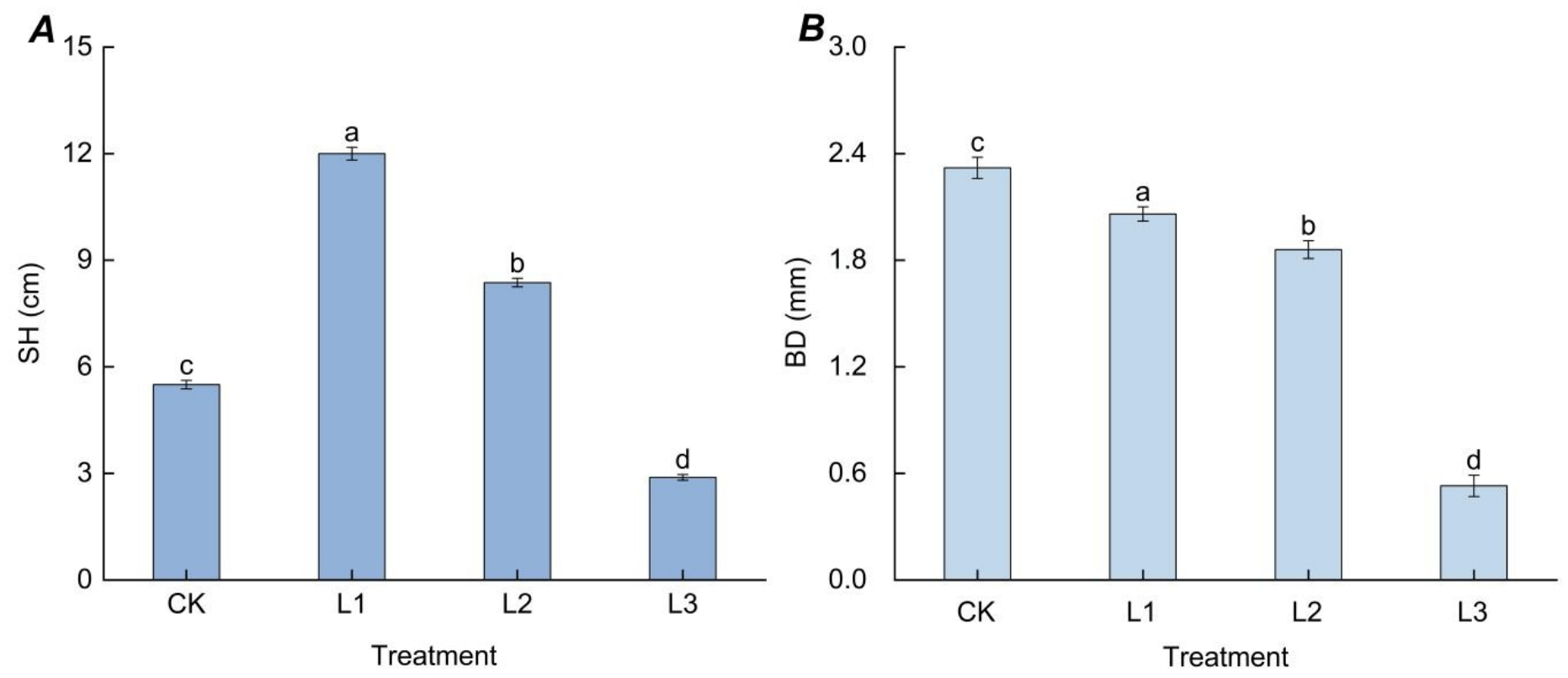
SH (*A*) and BD (*B*) of *S. sibirica* ‘Dong Hong’ under CK and nominal shading levels of L1, L2 and L3. Values are means ± SD of five seedlings per treatment. Different lowercase letters indicate differences among treatments within a panel (Tukey’s test, *P* < 0.05). SH, seedling height; BD, basal diameter; CK, full sunlight; L1, L2 and L3, nominal shading levels of 30%, 50% and 70%, respectively.

### Biomass and allocation

LB, SB, RB and TB showed consistent treatment rankings, with the highest values under L1, followed by CK, L2 and L3 (Fig. 2*A*). TB was 8.19 ± 0.24 g under L1 and 7.09 ± 0.26 g under CK, compared with 5.40 ± 0.08 g under L2 and 4.24 ± 0.07 g under L3. Despite these differences in absolute biomass, RBP remained similar across treatments, accounting for approximately 54-55% of TB (Fig. 2*B*). LBP was higher under L2 and L3 than under CK and L1, whereas SBP was lower under L2 and L3 than under CK. CK and L1 did not differ significantly in any biomass proportion. Thus, greater biomass under L1 coincided with an allocation pattern similar to CK, while the higher LBP under L2 and L3 was accompanied by lower absolute LB.

**Fig. 2.**
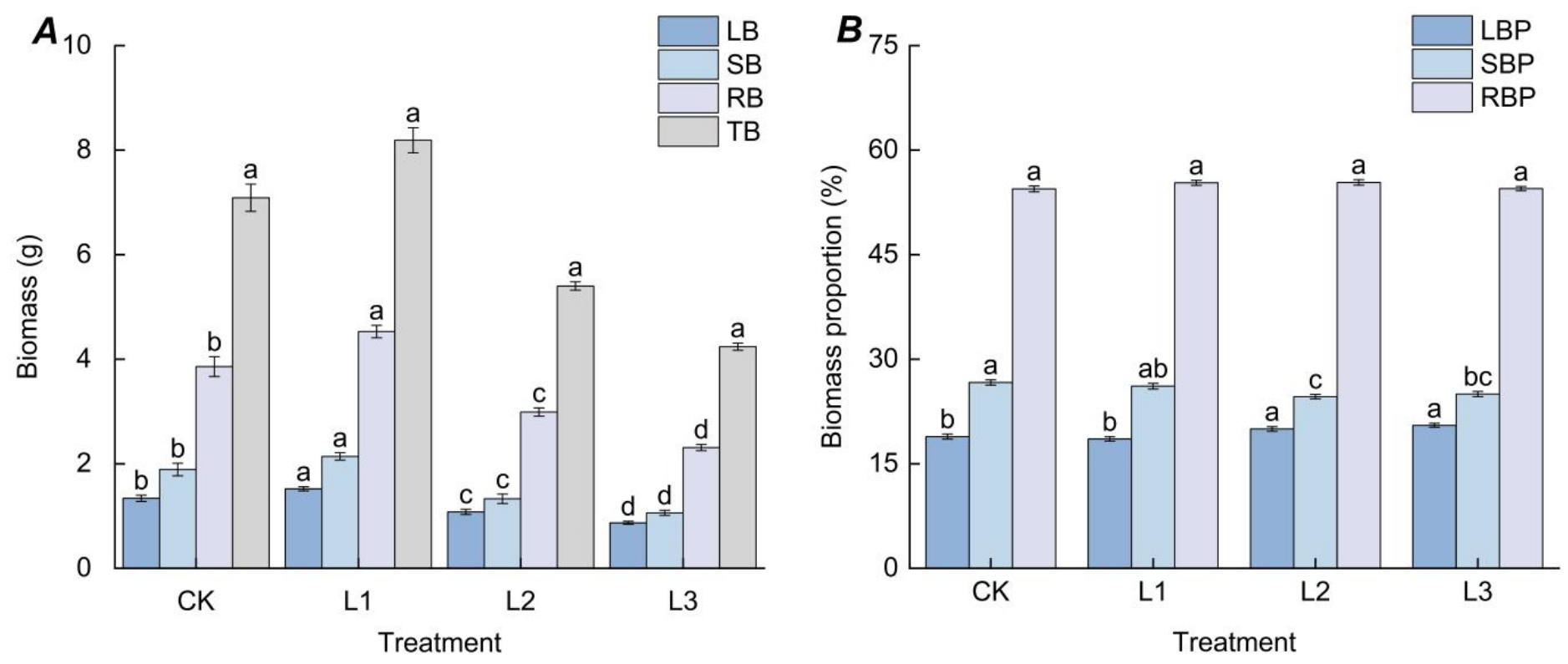
Organ and total dry biomass (*A*) and biomass allocation proportions (*B*) of *S. sibirica* ‘Dong Hong’ under CK and nominal shading levels of L1, L2 and L3. Values are means ± SD of five seedlings per treatment. Different lowercase letters indicate differences among treatments for each trait (Tukey’s test, *P* < 0.05). LB, leaf biomass; SB, stem biomass; RB, root biomass; TB, total biomass; LBP, leaf biomass proportion; RBP, root biomass proportion; SBP, stem biomass proportion; CK, full sunlight; L1, L2 and L3, nominal shading levels of 30%, 50% and 70%, respectively.

### Root morphology

Root morphological responses differed among traits and shading levels (Table 1). RSA, RAD and TRL did not differ significantly between L1 and CK, whereas NR and RV were lower under L1. Under L2, RSA and TRL were lower than under CK and L1, while RAD remained statistically similar across these three treatments. NR and RV did not differ significantly between L1 and L2. L3 had the lowest values for all five root traits, each significantly lower than in the other treatments. Thus, the similarity in TRL and RAD between L1 and CK was accompanied by differences in NR and RV, whereas L3 showed consistently lower values across all measured root traits. Representative whole-seedling photographs are shown in Fig. 3.

**Table 1.** Root morphology of *S. sibirica* ‘Dong Hong’ under CK and nominal shading levels of L1, L2 and L3. Values are means ± SD of five seedlings per treatment. Different lowercase letters within a column indicate differences among treatments (Tukey’s test, *P* < 0.05). RSA, root surface area; NR, root tip number; RAD, mean root diameter; RV, root volume; TRL, total root length; CK, full sunlight; L1, L2 and L3, nominal shading levels of 30%, 50% and 70%, respectively.

| Treatment | RSA (mm<sup>2</sup>) | NR (number) | RAD (mm) | RV (mm<sup>3</sup>) | TRL (mm) |
|---|---|---|---|---|---|
| CK | 74.92 ± 2.94<sup>a</sup> | 1 339.67 ± 87.14<sup>a</sup> | 0.39 ± 0.02<sup>a</sup> | 2.64 ± 0.25<sup>a</sup> | 533.88 ± 55.97<sup>a</sup> |
| L1 | 68.20 ± 1.88<sup>a</sup> | 1 119.00 ± 57.71<sup>b</sup> | 0.41 ± 0.05<sup>a</sup> | 1.57 ± 0.67<sup>b</sup> | 537.01 ± 6.61<sup>a</sup> |
| L2 | 50.57 ± 3.34<sup>b</sup> | 998.33 ± 56.37<sup>b</sup> | 0.36 ± 0.03<sup>a</sup> | 1.44 ± 0.20<sup>b</sup> | 406.34 ± 35.48<sup>b</sup> |
| L3 | 7.64 ± 0.67<sup>c</sup> | 215.00 ± 31.56<sup>c</sup> | 0.28 ± 0.01<sup>b</sup> | 0.12 ± 0.03<sup>c</sup> | 91.73 ± 3.84<sup>c</sup> |

**Fig. 3.**
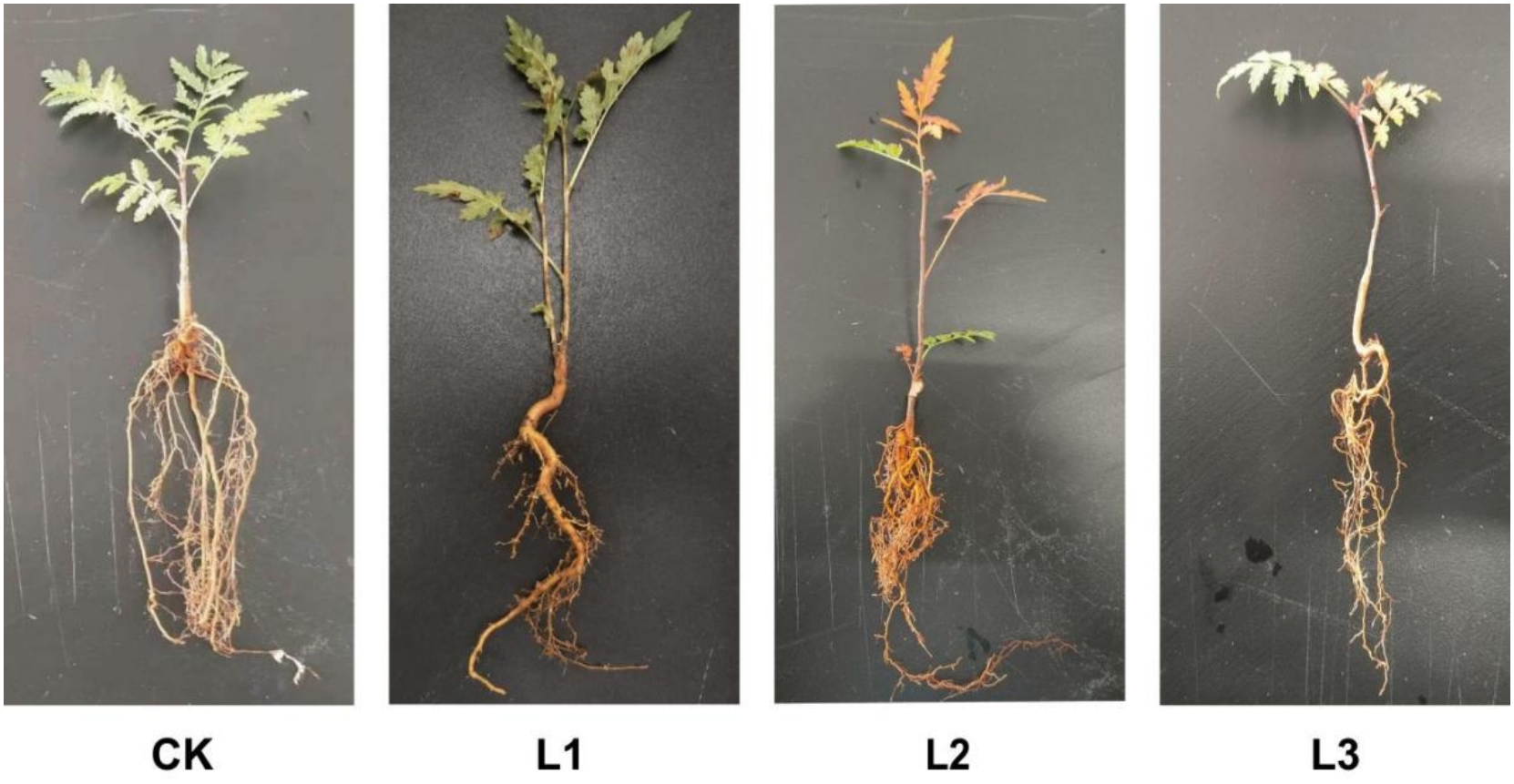
Representative seedlings of *S. sibirica* ‘Dong Hong’ under CK and nominal shading levels of L1, L2 and L3. Additional seedlings were sampled and photographed on 30 October 2024 for illustrative purposes. Photographs illustrate seedling appearance; quantitative root traits are reported in Table 1. Images are not intended for quantitative size comparisons. CK, full sunlight; L1, L2 and L3, nominal shading levels of 30%, 50% and 70%, respectively.

### Physiological responses

CHL content and SP content showed consistent treatment rankings, with the highest values under L1, followed by CK, L2 and L3 (Fig. 4*A, C*). Under L1, CHL content and SP content reached 8.39 ± 0.21 mg·g^-1^ FW and 0.6894 ± 0.012 mg·g^-1^ FW, respectively. In contrast, INV activity decreased progressively from 0.0061 ± 0.00012 U·g^-1^ FW under CK to 0.0031 ± 0.00007 U·g^-1^ FW under L3 (Fig. 4*B*). Thus, higher CHL content and SP content under L1 coincided with lower INV activity than under CK. Under L2 and L3, all three traits were lower than under CK, with the lowest values occurring under L3.

**Fig. 4.**
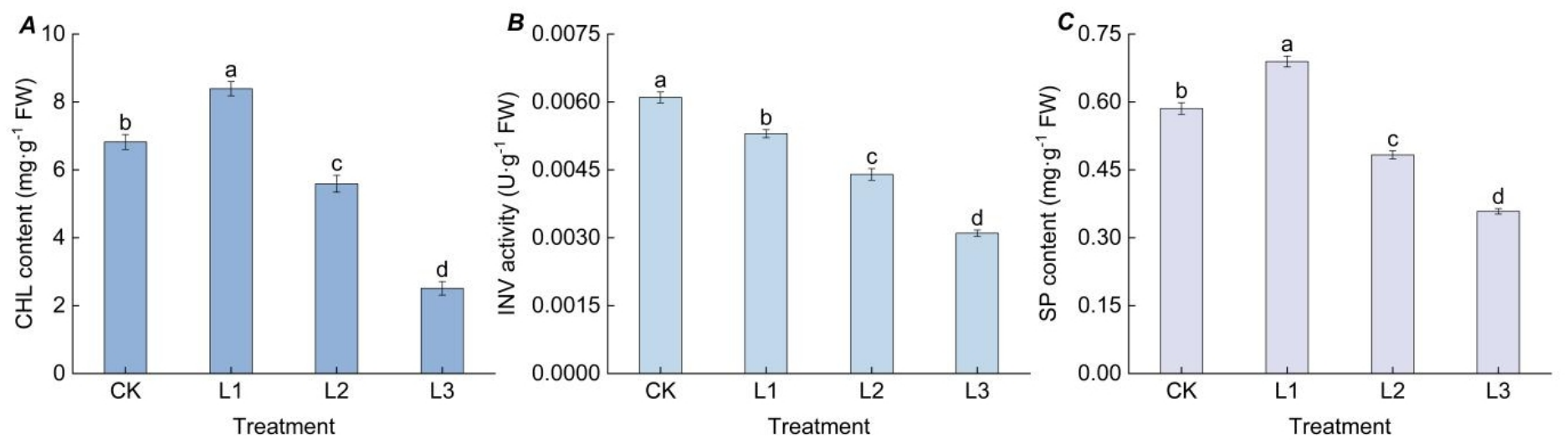
CHL content (*A*), INV activity (*B*) and SP content (*C*) of *S. sibirica* ‘Dong Hong’ under CK and nominal shading levels of L1, L2 and L3. Values are means ± SD of five seedlings per treatment. Different lowercase letters indicate differences among treatments within a panel (Tukey’s test, *P* < 0.05). CHL, chlorophyll; INV, invertase; SP, soluble protein; FW, fresh weight; CK, full sunlight; L1, L2 and L3, nominal shading levels of 30%, 50% and 70%, respectively.

### Leaf surface micromorphology

Stomatal dimensions and density showed different responses across treatments (Table 2). SL decreased progressively from CK to L3, whereas SA was highest under CK and lowest under L2, with a higher value under L3 than under L2. Both SL and SA were lower in all shaded treatments than in CK. In contrast, SDD did not differ significantly among treatments, indicating that the observed differences in stomatal dimensions were not accompanied by detectable differences in density. Qualitative observations showed a plate-like wax appearance under CK and L1 and a granular appearance under L2 and L3 (Fig. 5). These observations describe surface appearance rather than wax quantity or function.

**Table 2.** Stomatal traits and qualitative epicuticular wax appearance on the abaxial leaf surface of *S. sibirica* ‘Dong Hong’ under CK and nominal shading levels of L1, L2 and L3. Values are means ± SD of five seedlings per treatment. Different lowercase letters indicate differences among treatments within a column (Tukey’s test, *P* < 0.05). No significance test is implied for wax appearance. SL, stomatal length; SA, stomatal area; SDD, stomatal density; CK, full sunlight; L1, L2 and L3, nominal shading levels of 30%, 50% and 70%, respectively.

| Treatment | SL (µm) | SA (µm<sup>2</sup>) | SDD (stomata·mm<sup>-2</sup>) | Wax appearance |
|---|---|---|---|---|
| CK | 123.81 ± 5.23<sup>a</sup> | 1 934.88 ± 170.41<sup>a</sup> | 4.36 ± 0.17<sup>a</sup> | plate-like |
| L1 | 110.58 ± 7.52<sup>b</sup> | 1 236.72 ± 169.37<sup>b</sup> | 4.63 ± 0.23<sup>a</sup> | plate-like |
| L2 | 91.17 ± 13.07<sup>c</sup> | 647.80 ± 180.11<sup>d</sup> | 4.19 ± 0.11<sup>a</sup> | granular |
| L3 | 74.30 ± 4.20<sup>d</sup> | 767.79 ± 187.48<sup>c</sup> | 4.38 ± 0.09<sup>a</sup> | granular |

**Fig. 5.**
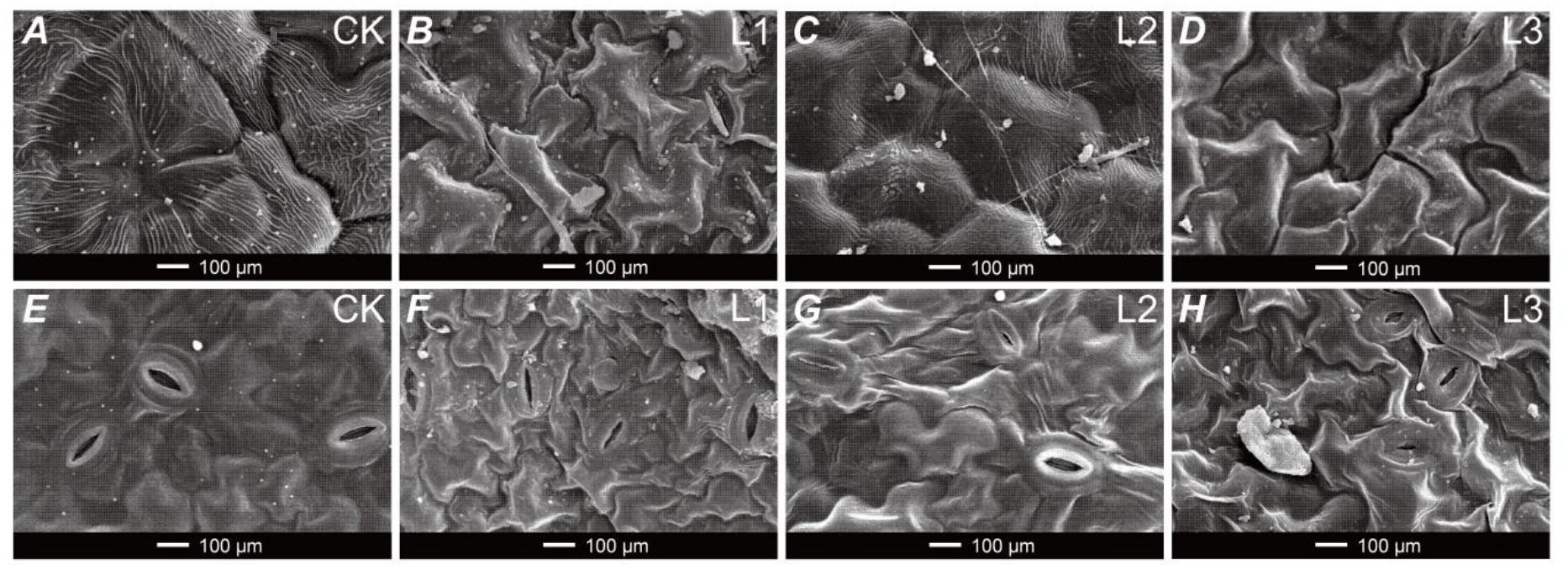
Scanning electron micrographs of the abaxial leaf surface of *S. sibirica* ‘Dong Hong’ under CK and nominal shading levels of L1, L2 and L3. *A–D*, qualitative epicuticular wax appearance under CK, L1, L2 and L3, respectively; *E–H*, stomatal morphology under the corresponding treatments. Scale bars = 100 µm. CK, full sunlight; L1, L2 and L3, nominal shading levels of 30%, 50% and 70%, respectively.

### Exploratory trait associations

Spearman correlations calculated from the four treatment means revealed shared rankings among several traits (Fig. 6). LB, SB, RB and TB were positively correlated with one another and with CHL content, SP content, RAD and TRL (ρ = 1.00). In contrast, LBP showed inverse rankings with these traits (ρ = ™1.00), indicating that a higher leaf mass fraction coincided with lower absolute biomass. BD, INV activity, RSA, NR, RV and SL also shared identical rankings (ρ = 1.00). SH was more closely associated with biomass, CHL content and SP content (ρ = 0.80) than with BD (ρ = 0.40). SDD showed comparatively weak associations with the other traits (|ρ| ≤ 0.40). These correlations describe covariation among treatment means rather than relationships among individual seedlings or causal effects.

**Fig. 6.**
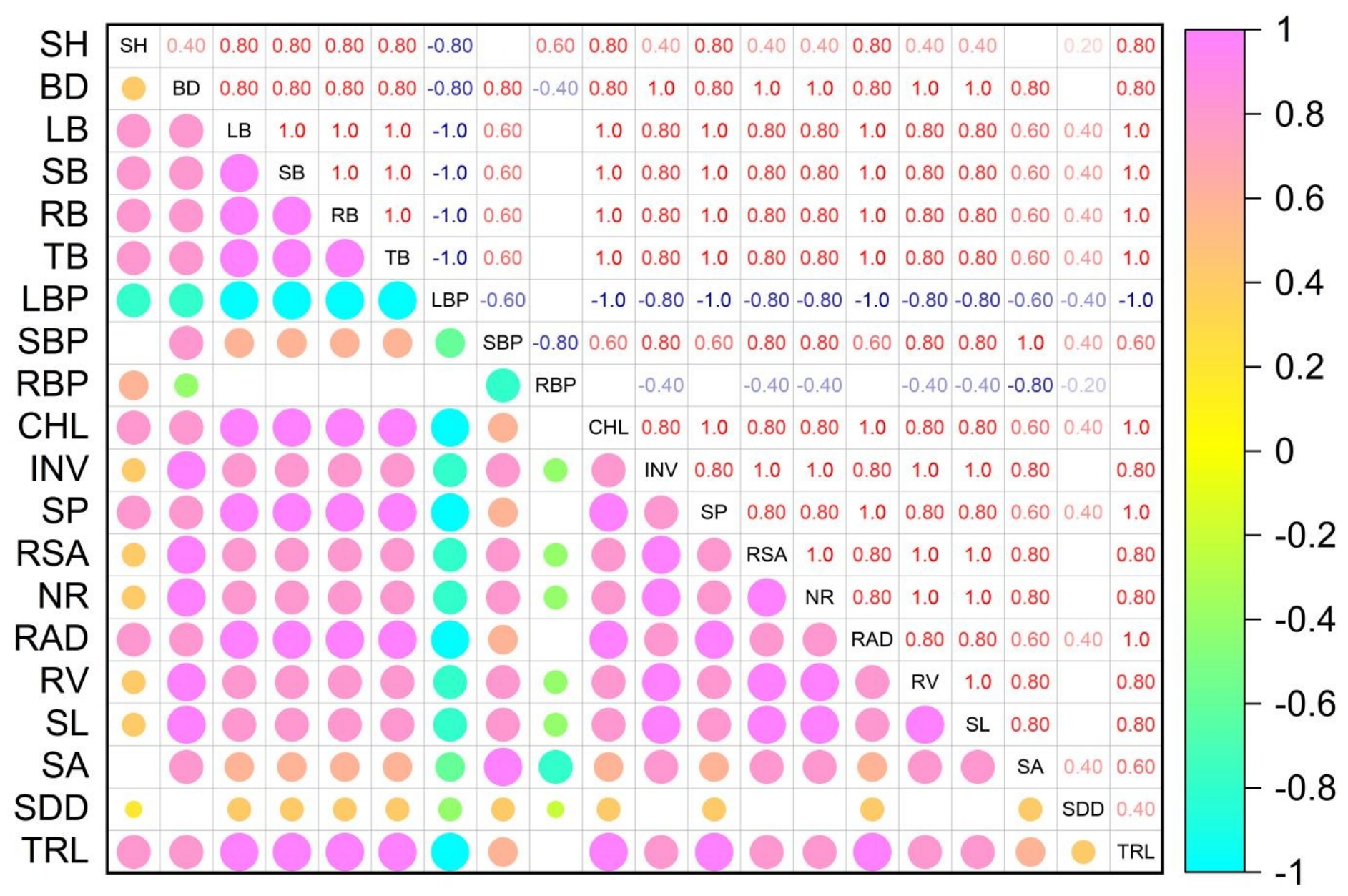
Exploratory Spearman rank correlations among growth, biomass allocation, physiological, root morphological and stomatal traits of *S. sibirica* ‘Dong Hong’. Correlations were calculated using the four treatment means (*n* = 4). The upper triangle shows Spearman correlation coefficients (ρ), and the lower triangle represents their direction and magnitude using circle color and size. Light magenta and cyan indicate positive and negative correlations, respectively; larger circles indicate stronger absolute correlations. The matrix describes covariation among treatment means and does not imply causal relationships. SH, seedling height; BD, basal diameter; LB, leaf biomass; SB, stem biomass; RB, root biomass; TB, total biomass; LBP, leaf biomass proportion; SBP, stem biomass proportion; RBP, root biomass proportion; CHL, chlorophyll; INV, invertase; SP, soluble protein; RSA, root surface area; NR, root tip number; RAD, mean root diameter; RV, root volume; SL, stomatal length; SA, stomatal area; SDD, stomatal density; TRL, total root length.

## Discussion

The responses of *S. sibirica* ‘Dong Hong’ to shading differed among traits, indicating that nursery performance cannot be assessed from SH alone. L1 had the highest SH and biomass, with TB reaching 8.19 ± 0.24 g compared with 7.09 ± 0.26 g under CK. Thus, greater SH under L1 coincided with greater final dry mass rather than elongation alone. However, BD remained lower than under CK, revealing a difference between biomass production and stem thickening. Under L2, taller seedlings had lower biomass and BD than CK, while L3 had the lowest values for all three traits. These patterns identify a potential benefit of L1 under the experimental conditions. The taller but thinner stems under L1 and L2 are consistent with altered shoot development under reduced light. Such morphology may favour light interception, but its implications for mechanical stability and subsequent establishment remain uncertain (Damtew et al., 2025; Syahputra et al., 2026). Greater SB under L1 did not correspond to greater BD, suggesting that stem mass and dimensions responded differently. Moreover, the experiment assessed final size rather than growth trajectories. The results therefore support evaluating SH, BD and biomass together and avoiding the assumption that the tallest seedlings necessarily provide the best planting material (Medeiros, 2017; Zhou et al., 2025).

CHL content and SP content followed the same treatment ranking as biomass, with maxima under L1. Higher CHL content could reflect adjustment of the pigment pool to reduced irradiance, potentially supporting light capture (Liu et al., 2026; Zhang et al., 2025). The higher SP content under L1 indicates a larger soluble protein pool per unit leaf FW, but does not identify the proteins or metabolic pathways involved. Partial shading may also have reduced excessive radiation or thermal loads while maintaining sufficient light for growth (Guan et al., 2024). However, photosynthetic gas exchange, CHL fluorescence and leaf temperature were not measured; therefore, reduced photoinhibition or thermal stress cannot be confirmed. A previous study reported favourable growth and physiological responses to intermediate shading in *Melaleuca alternifolia*, supporting this possibility in other nursery systems (Li et al., 2025). The lower CHL content, SP content and biomass values under L2 and L3 indicate that stronger shading did not provide progressively greater benefits. Reduced light availability may have constrained carbon acquisition, while changes in leaf development or senescence could also have contributed to the physiological differences (Lv et al., 2025; Sha et al., 2026). INV activity declined across the shading gradient despite the biomass maximum under L1, further demonstrating that individual biochemical indicators did not uniformly track whole-plant performance (Lou et al., 2026). Although INV activity can influence growth through sucrose metabolism, its effects depend on tissue and isoform (Galibina et al., 2019).

Biomass allocation provides additional context for these responses. CK and L1 had similar leaf, stem and root mass fractions despite their different TB, indicating that greater biomass under L1 occurred without a pronounced change in proportional allocation. Conversely, L2 and L3 had higher leaf mass fractions but lower absolute LB. This pattern may represent relative maintenance of foliage as plant mass declined, rather than increased leaf production (Sevillano et al., 2016). RBP remained approximately 54–55% across treatments, whereas root morphological responses were less uniform. Under L1, TRL and RAD were comparable to CK, but NR and RV were lower. Similar TRL therefore did not imply equivalent branching or root system organisation. The higher root dry mass under L1 also indicates that dry mass and scanned morphological traits describe different aspects of root development. Their relationship requires examination of root diameter classes and tissue properties before functional conclusions can be drawn (Dong et al., 2022). Under L3, consistently low root morphological values accompanied reduced biomass, suggesting coordinated limitations to shoot and root development. Stomatal dimensions responded more consistently than density. SL declined with increasing shade, whereas SA was lowest under L2. SDD was numerically highest under L1, but SDD did not differ significantly among treatments. Smaller stomata have been associated with faster responses in comparative studies (Drake et al., 2013), but stomatal response kinetics and conductance were not measured here. The smaller stomata under L2 and L3 also coincided with lower biomass, indicating that reduced stomatal dimensions alone did not imply improved growth. Qualitative differences in surface appearance were observed between CK–L1 and L2–L3. Light-dependent changes in epidermal wax deposition have been reported in *Hosta* (Guan et al., 2024), but chemical analysis and permeability measurements would be needed to establish the functional significance of the present observations.

Comparisons with other Sorbus species emphasise the conditional nature of shading responses. A previous study reported favourable performance of *S. tianschanica* at 30% of full sunlight (Chen et al., 2025b). This nominal exposure corresponds to L3 here, which had the lowest biomass. Differences in species, seedling age and cultivation environment may contribute to these contrasting outcomes. Furthermore, the present experiment covered only 20 July to 28 October 2024. Seasonal variation documented in previous reports on *Acer miaotaiense* subsp. *yangjuechi* likewise supports considering treatment timing alongside shading intensity (Chen et al., 2025c). Across the sampled dates and times, mean illuminance was 18 520 lx under CK and 12 601, 9 136 and 5 640 lx under L1, L2 and L3, respectively. However, measurements on three clear-sky days did not characterise cumulative light exposure over the 100-day treatment period or establish whether light conditions were typical of the region. Consequently, comparisons with other studies should distinguish nominal shading levels from measured light conditions (Wang et al., 2024). Container conditions further constrain application. Limited rooting space can influence growth and physiology, while manual watering and the absence of fertilisation define the cultivation context of this study. Responses may differ in larger containers or field soils (Liu et al., 2023). These limitations are particularly relevant when considering transfer to nurseries with different climates, substrates or irrigation practices.

Finally, correlations based on four treatment means describe shared rankings rather than independent physiological mechanisms, and associations between TB and its components partly reflect mathematical dependence. Overall, L1 warrants further evaluation as a conditional nursery option, while L3 offered no growth advantage during this study. These findings do not establish a year-round shading prescription. Future trials should compare seasonal shading schedules under measured light and moisture conditions and assess stem development, root function, overwintering and post-transplant establishment. Such validation would determine whether the observed biomass benefit translates into improved nursery and landscape performance.

## Conclusion

*S. sibirica* ‘Dong Hong’ seedlings showed trait-dependent responses to shading during the 100-day summer-to-autumn pot experiment. L1 was associated with the highest SH, biomass, CHL content and SP content, but BD, NR and RV remained lower than under CK. L2 and L3 had lower biomass than CK, with L3 showing the lowest values across all measured root morphological traits. These findings identify nominal 30% shading as a potential option for further evaluation in comparable container nurseries, rather than a universally optimal regime. Its suitability depends on production objectives, seasonal light availability and cultivation conditions. Restricted rooting volume, limited environmental monitoring and the absence of multi-season and field validation constrain broader application. Shading intensity and duration should therefore be evaluated locally, with attention to stem development, root performance, overwintering and establishment after transplanting.

## Abbreviations

ANOVA: analysis of variance
BD: basal diameter
CHL: chlorophyll
CK: full sunlight
DNS: 3,5- dinitrosalicylic acid
EDTA: ethylenediaminetetraacetic acid
FW: fresh weight
INV: invertase
LB: leaf biomass
LBP: leaf biomass proportion
NR: root tip number
PVPP: polyvinylpolypyrrolidone
Q-Q: quantile-quantile
RAD: mean root diameter
RB: root biomass
RBP: root biomass proportion
RSA: root surface area
RV: root volume
SA: stomatal area
SB: stem biomass
SBP: stem biomass proportion
SD: standard deviation
SDD: stomatal density
SEM: scanning electron microscopy
SH: seedling height
SL: stomatal length
SP: soluble protein
TB: total biomass
TRL: total root length.

## Acknowledgments

The authors sincerely thank the College of Life Science, Agriculture and Forestry of Qiqihar University for providing technical support and experimental facilities in this study.

## Notes

*Data Availability Statement:* The data supporting the findings of this study are available from the corresponding author upon reasonable request.

### Competing Interest Statement

The authors have declared no competing interest.

### Summary of Updates

The Abstract and Introduction now clarify the research objectives, cultivar background, ornamental characteristics and relevance to container nursery management. The study is framed as an assessment of trait-specific responses rather than an attempt to establish a universally optimal shading regime. The Materials and methods section has been expanded to specify treatment dates, cultivation conditions, sampling procedures, biomass drying criteria, root scanning, physiological assays and scanning electron microscopy. Light measurements are correctly reported as illuminance rather than photosynthetic photon flux density, with measured values and relative attenuation distinguished from nominal shading levels. Statistical procedures and the averaging of technical measurements are also clarified. An error in biomass sample aggregation was corrected, changing the reported comparison between L1 and CK; the corresponding results, figures and interpretations have been updated consistently. The Results now emphasise treatment rankings, similarities and contrasts among traits, with fewer percentage comparisons. Stomatal density is described as not differing significantly among treatments, and wax appearance is interpreted qualitatively. Spearman correlations based on four treatment means are explicitly presented as exploratory descriptions without significance testing or causal inference. The Discussion has been rewritten around the experimental findings, particularly the greater biomass but reduced stem thickness and altered root morphology under L1. Possible mechanisms are distinguished from directly measured responses, and the absence of gas-exchange, fluorescence and leaf-temperature measurements is acknowledged. The Conclusion now presents 30% shading as a conditional option requiring further evaluation, with explicit limitations arising from restricted rooting volume, limited environmental monitoring and a single summer-to-autumn experiment. Abbreviations, units, captions and figure presentation have been revised, and highlights and a graphical abstract have been prepared.

